# Naming Human Diseases: Ethical Principles of Curating Exclusive Substitute for Inopportune Nosology

**DOI:** 10.1101/2021.05.01.442270

**Authors:** Zhiwen Hu, Ya Chen, Yuling Song, Zhongliang Yang, Hui Huang

## Abstract

**Background:** In the medical sphere, understanding naming conventions strengthen the integrity of naming human diseases remains nominal rather than substantial yet. Since the current nosology-based standard for human diseases could not offer a one-size-fits-all corrective mechanism, many idiomatic but flawed names frequently appear in scientific literature and news outlets at the cost of sociocultural impacts.

**Objective:** We attempt to examine the ethical oversights of current naming practices and propose heuristic rationales and approaches to determine a pithy name instead of an inopportune nosology.

**Methods:** First, we examined the compiled global online news volumes and emotional tones on some inopportune nosology like *German measles, Middle Eastern Respiratory Syndrome, Spanish flu, Hong Kong flu*, and *Huntington’s disease* in the wake of COVID-19. Second, we prototypically scrutinize the lexical dynamics and pathological differentials of *German measles* and common synonyms by leveraging the capacity of the Google Books Ngram Corpus. Third, we demonstrated the empirical approaches to curate an exclusive substitute for an anachronistic nosology *German measles* based on deep learning models and *post-hoc* explanations.

**Results:** The infodemiological study shows that the public informed the offensive names with extremely negative tones in textual and visual narratives. The findings of the historiographical study indicate that many synonyms of *German measles* did not survive, while *German measles* became an anachronistic usage, and *rubella* has taken the dominant place since 1994. The PubMedBERT model could identify *rubella* as a potential substitution for *German measles* with the highest semantic similarity. The results of the semantic drift experiments further indicate that *rubella* tends to survive during the ebb and flow of semantic drift.

**Conclusions:** Our findings indicate that the nosological evolution of anachronistic names could result in sociocultural impacts without a corrective mechanism. To mitigate such impacts, we introduce some ethical principles for formulating an improved naming scheme. Based on deep learning models and *post-hoc* explanations, our illustrated experiments could provide hallmark references to the remedial mechanism of naming practices and pertinent credit allocations.

## Introduction

### Background

Terminology is the crystallization of human scientific and technological knowledge in natural language. In the medical sphere, appropriate names were deliberately invented for the designation of human diseases with pathological characteristics. However, underrepresented emphasis has been placed on the nomenclature of human diseases. The current wave of destigmatization calls for constant introspection of the offensive appellations of human diseases [1–3]. In the same week, the anachronistic usage of *German measles* in the leading journals *Nature* and *Science* without any caution implies that some strongly-held but flawed names may brand social stigma and discrimination [4–6].

In the 19^th^ century, the name *rubella* was proposed as a substitute for German term *rötheln*, then the epidemic neologism *German measles* was gradually accepted as idiomatic usages [7–16]. However, anachronistic usages like that violate the latest naming protocols of the World Health Organization (WHO) – stigmatizing a specific country and its residents [1]. Arguably, the looming worry is to reignite the torch of discrimination and fuel the current infodemic unconsciously [3,17–20].

### Study Objectives

Based on extensive literature review, this study aims to punctuate heuristic introspection of naming practices for human diseases and address the following research issues:

1. Did the anachronistic names like *German measles* cost social impacts?
2. What are the diachronic discourses of *German measles* and common synonyms? What can we learn from the lexical evolution?
3. Should we hash out inopportune names like *German Measles*? And How?
4. What are the pertinent principles of curating the exclusive substitute for an anachronistic nosology?

## Methods

Rich collections of the printed or digital imprint of social individuals are formidable proxies to determine the dynamic pragmatics patterns of practical utterances and reveal the collective human behaviours from sociocultural preferences [21,22]. Following the Preferred Reporting Items for Systematic Reviews and Meta-Analyses (PRISMA) guidelines [23], here we orchestrate rich metadata available to unveil the scientific paradigms via the following experiments (**Multimedia Appendix 1**).

### Infodemiological study

In the global online news coverage experiments, we aim to unveil the scientific paradigms of the diachronic discourse and emotional tone. Here, the metadata analysis aims to demonstrate the emotional polarity of the public in the context of global online news on *German measles, Middle Eastern Respiratory Syndrome, Spanish flu, Hong Kong flu*, and *Huntington’s disease* over time, respectively.

First, the code scheme was curated following three main principles that we established before [24]. According to the code scheme, the search formulas are available in **Multimedia Appendix 2**. Second, the unbiased and comprehensive metadata of global online news coverage and emotional tone retrieved through the open project GDELT Summary between December 30, 2019 (the outbreak of COVID-19) and May 8, 2021 (the Sixth anniversary of *World Health Organization Best Practices for the Naming of New Human Infectious Diseases*), including the textual and visual narratives of different queries [25,26]. Finally, by leveraging the capacity of GDELT’s machine translate and neural network image recognition [26], the instant news portfolio in **Figure 1** summarizes the textual and visual narratives of different queries in 65 multilingual online news. The volume ratio is the total volume of matching articles divided by the total number of all articles monitored by GDELT. The emotional tone is the average tone of all matching documents, and the normalized score ranges from −10 (extremely negative) to +10 (extremely positive) based on the tonal algorithm.

**Figure 1.**
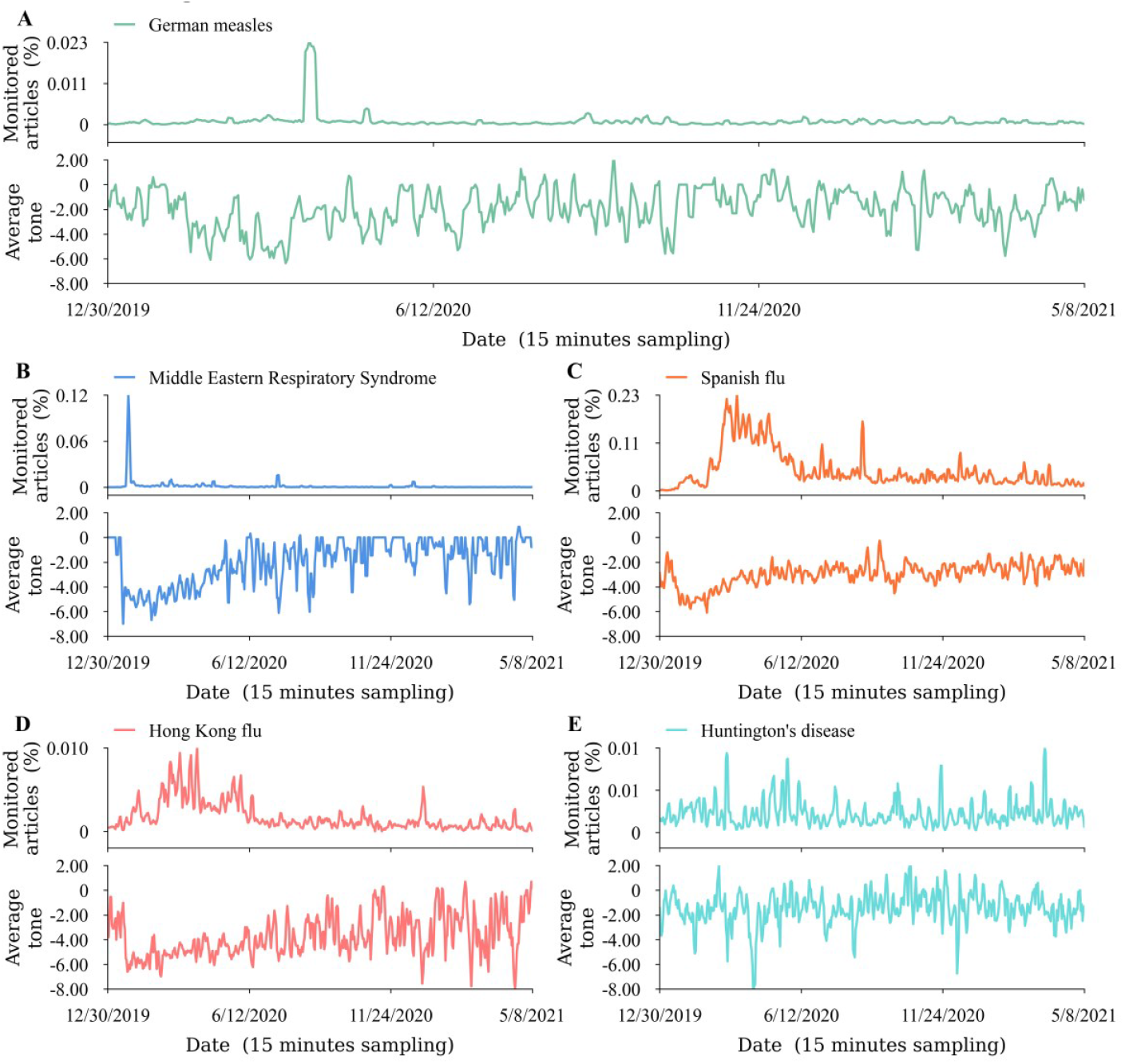
Prevailing stereotypes of stigmatizing names with negative tones in the wake of COVID-19 pandemic. The global instant news portfolio on GDELT Summary summarizes the textual and visual narratives of different queries in 65 multilingual online news: A, *German measles*; B, *Middle Eastern Respiratory Syndrome*; C, *Spanish flu*; D, *Hong Kong flu*; and E, *Huntington’s disease*. The upper panels display the percent of all global online news coverage over time. The lower panels show the average emotional tone of all news coverage from extremely negative to extremely positive. The temporal resolution of sampling is 15 minutes per day.

### Historiographical study

The Google Books Ngram Corpus (GBNC) is a unique linguistic landscape that benefits from centuries of development of rich grammatical and lexical resources as well as its cultural context [27]. It contains *n*-grams from approximately 8 million books, or 6% of all books published in English, Hebrew, French, German, Spanish, Russian, Italian, and Chinese. The GBNC covers data logs from 1500 to 2019. A unigram (1-gram) is a string of characters uninterrupted by a space, and an *n*-gram (*n* consecutive words) is a sequence of a 1-gram, such as *morbilli* (unigram), *rubeola* (unigram), *rubella* (unigram), *Rötheln* (unigram), and *German measles* (bigram). In this study, by retrieving the use frequency of a specific lexicon in historical development, we first obtain a glimpse of the nature of historical evolution in **Figure 3**.

Then, as we continue to stockpile seminal patterns in **Figure 3**, some have argued that correlation is threatening to unseat causation as the bedrock of scientific storytelling before. We must punctuate heuristic cautions of wrestling with information from retrospective sources, cross-validation, and the reassembly of the whole story. Finally, we provide compelling arguments to the extent of understanding the underneath nature of lexical dynamics and pathological differentials based on authentic materials and critical examination.

### Semantic similarity experiments

Based on the epistemic results of the above historiographical study, as an exemplificative case, we could construct the initial candidates of *German measles*, which includes *morbilli, rubeola, rubella*, and *rötheln*. Relatedly, as prior knowledge, the term *rotheln* is ordinarily used as a translation of the German term *rötheln* in literature. From the outset, it’s reasonable to expand the initial candidates to *morbilli, rubeola, rubella, rötheln*, and *rotheln*.

Directed at five expanded candidate words, we employed the BERT model and PubMedBERT model to quantify the semantic similarities between them, respectively. The cosine similarity formulas to calculate semantic relevance is as follows:

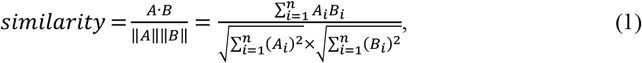

where *A* and *B* denote two ectors, *A*_*i*_ and *B*_*i*_(*i*= 1 … *n*) represent the components of vector *A* and *B*.

The BERT model and PubMedBERT model have the same architecture with different corpora for preliminary training and pre-training (**Figure 4**). Coupling with a multi-layer bidirectional transformer encoder and bidirectional self-attention, the BERT and PubMedBERT models are more sensitive to semantics than the constrained self-attention used by GPT-2 model. The former uses the BookCorpus (800M words) and English Wikipedia (2,500M words) for training, its multilingual pre-training model can handle over more than 100 languages [28]. The latter model uses the latest collection of PubMed abstracts (14M abstracts, 3.2B words, 21GB), and its pre-training model can facilitate understanding the word semantic in the medical field [29]. The two models used the Wordpiece embeddings with their own token vocabularies. The two models are capable to verify the homology between *rötheln*, and *rotheln*, and identify the target word with the closest similarity to *German measles* in the initial candidates (**Figure 5**). For *post-hoc* explanations, PubMedBERT-generated case studies available facilitate to demystify the typical scenarios in pre-training narratives.

### Semantic drift experiments

We analyzed the dynamic evolution of the five keywords *German measles, morbilli, rubeola, rubella*, and *rötheln*. In the experiments, the two-stage text corpora were retrieved from GBNC. Specifically, we choose two time periods: one or two hundred years after the word appeared and 1950 to 2020. Each keyword has two corpora, and each corpus has 1,000 targeted snippets published in two time periods with random sampling.

To accurately demonstrate the semantic evolution of each keyword, we orchestrate their synchronic and diachronic semantic structures. Since a word’s historical meaning can be inferred by its most semantically similar words in two time periods, we could track down how the semantics of words change over time [30,31]. Firstly, we used word co-occurrence matrix to build semantic representations in two time periods, in this way the meaning of the word could be approximated in the contexts over time. By the word co-occurrence matrix, we curated some semantic neighbors for the keywords *German measles, morbilli, rubeola, rubella*, and *rötheln*, respectively. Secondly, based on the word co-occurrence matrix, positive pointwise mutual information (PPMI) matrix entries are given by:

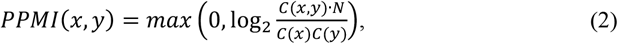

where *C* represents word co-occurrence matrix, *C*(*x, y*) refers to the number of co-occurrences of the words *x* and *y, C*(*x*) represents the number of occurrences of the word *x, C*(*y*) represents the number of occurrences of the word *y, N* represents the number of words in the corpus. Thirdly, we used singular value decomposition (SVD) to obtain a 4500×4500 matrix for the corpus of the two time periods. Finally, in **Figure 6**, we employed principal component analysis (PCA) to reduce the dimensions of word embeddings from 4,500 to 2 and then projected the uncharted latent patterns in the five word-embeddings clusters.

## Results

### Ethical oversights of naming practices

May 8, 2021 marks 6 years since the first best practices of new human infectious diseases was announced by WHO [1]. In recent years, we have witnessed many outbreaks of human diseases, with proper names given by stakeholders. Sometimes, diseases are initially given interim names or common names. Then, the proper names are officially ratified by the International Classification of Diseases (ICD) of WHO. Even so, each round of naming practice is not always successful [1,32,33]. Of them, *Middle Eastern Respiratory Syndrome* (MERS) [34], *Spanish flu* [35,36], *Hong Kong flu* (1968-1969) [37–39], and *Huntington’s disease* [40–43] have been accused of unnecessary social impacts in previous studies (**Figure 1**).

Naming conventions are not merely for naming diseases but for the vitality of science and the promotion of social progress [2,33,44,45]. Evidently, as shown in **Figure 1**, the results of the infodemiological study show that the global news outlets (in 65 languages) enjoy long-standing but flawed naming conventions with extremely negative tones, such as *German measles, Middle Eastern Respiratory Syndrome, Spanish flu, Hong Kong flu*, and *Huntington’s disease*. Admittedly, the coverage of affective tones is much negative than the standard portrayal assumes on average [46]. This finding highlights that these controversial stereotypes confounded the generally accepted norms at the cost of social progress, such as worsening acute stress of patients, provoking the backlash against particular communities, triggering unjustified slaughtering of food animals, and creating needless travel barriers or trade barriers [1–3,47,48].

Understanding how naming conventions strengthen the integrity of naming practices remains nominal rather than substantial yet. In the COVID-19 infodemic, multifarious monikers have become explicit consideration in the COVID-19 paper tsunami, and the global profusion of tangled hashtags has found their ways in daily communication [49]. Just as the remarks of the editorial of *Nature*, “As well as naming the illness, the WHO was implicitly sending a reminder to those who had erroneously been associating the virus with Wuhan and with China in their news coverage – including *Nature*. That we did so was an error on our part, for which we take responsibility and apologize.”[50] Unfortunately, many more stigmatized names somewhat aggravate the collective perceptual biases and contribute to recent backlash against Asians and diaspora [24,51]. Accordingly, scientists must verse themselves in naming conventions rather than feeding the trolls of stigma and discrimination.

### Ethical principles of naming human diseases

Of similar concern, we witness that many anachronistic names, from *Spanish flu* to *Zika*, and from *Lyme* to *Ebola*, are named after geographic places in our daily communications. But they are stigmatized cases and plain inaccurate (**Table 1**).

**Table 1.** Many diseases’ names have complained of both stigmatization and inaccurate.

| Name | Complains | References |
| --- | --- | --- |
| *Spanish Flu* | The 1918-19 influenza outbreak now often referred to as the *Spanish flu* did not even originate in Spain. | [52,53] |
| *Zika virus disease* | A mosquito-borne disease caused by *Zika virus* was first identified in Uganda in 1947 in monkeys, and later identified in humans in 1952. *Zika virus disease* was named after the Zika forest of Uganda. | [54] |
| *Lyme disease* | *Lyme disease* was named after the “original location”, the town of Old Lyme, Connecticut in 1975. More than 130 years ago, a German physician Alfred Buchwald first discovered the erythema migrans of what is now known to be *Lyme disease*. | [53] |
| *Ebola disease* | The hemorrhagic fever caused by the filovirus is named after the Ebola River at Legbala, Congo. Yambuku, a town situated 100 kilometers away from Legbala, was the first epicenter in 1976. | [53–55] |

In retrospect, WHO released the latest naming protocols of newly identified infectious diseases in 2015, as a supplement to the Reference Guide on the Content Model of the ICD-11 alpha drafted by ICD in 2011 [1,56,57]. In May 2019, the World Health Assembly (WHA) formally adopted the 11^th^ revision of the International Classification of Diseases (ICD-11). Specifically, we could crystallize the current recommendations into five protocols: avoidance of geographic locations (e.g., countries, cities, regions) ; avoidance of people’s names; avoidance of species/class of animal or food; avoidance of cultural, population, industry, or occupational references; and avoidance of terms that incite undue fear.

In theory, all Member States should follow the nosology-based standard to name a newly identified human disease at a later stage. On one hand, the global response has not always been smooth in practice. On the other hand, the international framework could not offer a one-size-fits-all surveillance mechanism for new designations and the pre-existed names. Currently, many inopportune names are widely professed in both scientific literature and news outlets without any caution, such as *Ebola, Rift Valley fever, Japanese encephalitis, Crimean Congo hemorrhagic fever, Chagas disease, Athlete’s foot, miner’s asthma, Marburg disease, Legionnaire’ disease, Creutzfeldt-Jakob disease, monkey pox, bird flu, equine encephalitis, paralytic shellfish poisoning, swine flu*, and so on [1,32,33]. Accordingly, after the swine flu of 2009 outbreak, some countries still banned pork imports, although *swine flu* cannot be transmitted from pigs at all. Moreover, they are scapegoating on the resurgence of stigma in the wake of geopolitical tensions and backlashes [53].

Still, the most pressing challenges concerning the nomenclature of human diseases depend on well-posed questions, including but not limited to: Who did coin a term to the designation of a specific disease? Is it still an appropriate name today? How many inopportune names garner in textbooks without any caution? How to map out an exclusive substitute for a flawed name? To that end, as a supplement to the current nosology-based nomenclature standard, we propose the ethical principles for naming a new human disease or renaming pre-existed nosology, as well as pertinent credit allocation (**Figure 2**).

**Figure 2.**
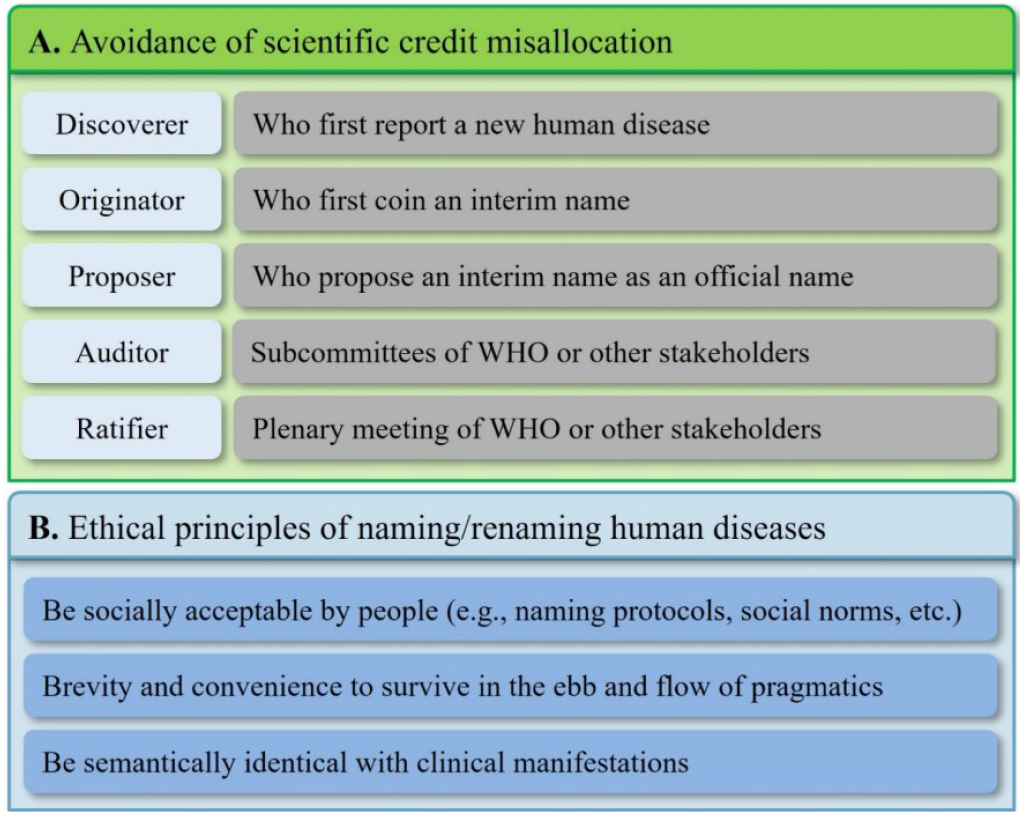
Proposal of ethical principles for the latest naming protocols of human diseases.

First, as shown in **Figure 2**, many contributors were involved in the general taxonomy and nomenclature process of human diseases, including the discover(s), originator(s), proposer(s), auditor(s), and ratifier(s). Without moral discernment, the Matthew effect of credit misallocations always discourages individual engagement in such practices [58]. Scientists, who preluded the accession of a particular disease, do not always earn their *bona fide* niches because of credit misallocation in the scientific narratives. Typically, the unsung originator Dick Thompson first coined the term “*Severe Acute Respiratory Syndrome*” (*SARS*) on 15 March 2003 [59]. His tour-de-force contribution is portrayed as a trivial anecdote. Similarly, Dr. Jean-Jacques Muyembe-Tamfum, one of the discoverers of *Ebola disease*, has been unsung until 2015 [60–62]. Thus, figuring out the seminal motivations of nomenclatures in routine obscurity generally presupposed that we could track down the original records [63,64]. To corroborate continuous introspection of previous multifarious findings, historians always find themselves buried in unending retrieval of tangled contingencies to pinpoint such inherent affiliations in pithy evidence.

Second, any proposed name of diseases should balance science, policy, and communication. Human diseases are often given names by stakeholders outside of the medical sphere. As a counterexample, *Pneumonoultramicroscopicsilicovolcanoconiosis*, referred to as pneumoconiosis or silicosis, was coined by Everett M. Smith in 1935 [65,66]. Literally, the 45-letter neologism is “a form of a chronic lung disease caused by the inhalation of fine silicate or quartz dust”, according to the *Oxford English Dictionary Online* (OED Online). Such a long disease entity does not roll off the tongue for efficient communication, and we should discard similar designations in scientific literature.

Third, erasing the stigmas of human diseases is a major public health priority. Whether name change will ever destigmatize a human disease or not is a cliché. As a case in point, *Schizophrenia* (also known as Kraepelin’s disease or Bleuler’s syndrome) was first adopted in 1937 as a translation of the German name *Schizophrenie*. To erase the potential stigma of *Schizophrenia*, the National Federation of Families with Mentally III in Japan requested the Japanese Society of Psychiatry and Neurology to replace the official term *Seishin Bunretsu Byo* with *Togo-Shicchou-Sho* in 2002 [67–69]. In the same vein, South Korea changed their official term *jeongshin-bunyeol-byung* to *johyun-byung* in 2011 [70]. Subsequently, lexical harm reduction became a bone of contention [68,71–75]. Keep the twists and turns of naming practices in mind, any novel proposal should go far beyond a mere semantic equivalent, as well as across cultures and languages. In practice, an alphanumeric code or a Greek letter is occasionally proposed for naming a pathogen or disease, but the opposers underline that such designations often introduce new confusion. For example, “filovirus-associated haemorrhagic fever 1” and “filovirus-associated haemorrhagic fever 2” were proposed for potential candidates instead of *Marburg disease* and *Ebola disease*, respectively [32]. Similarly, some Greek letters sound alike when they are translated into other languages, such as eta (the seventh letter of the Greek alphabet) and theta (the eighth letter of the Greek alphabet) [76]. With unreached consensus, some problematic notions are barning in our textbooks to educate generation after generation without any caution. Nonetheless, reassigning a curated standard name is the corrective approach to destigmatize a flawed name of infectious diseases or noncommunicable diseases.

Last but not at least, sociocultural costs of inappropriate disease names receive little attention [53]. Scientists mostly work to reduce the physical toll of diseases, although many of them endorse that disease names could be problematic [32]. Consistent with our findings, in most instances, inappropriate disease names remain in our blind spot, partially because the lasting damage they cause is difficult to quantify. Just as the remarks of Dr. Keiji Fukuda, Assistant Director-General for Health Security of the WHO, “this may seem like a trivial issue to some, but disease names really do matter to the people who are directly affected.”[48] Admittedly, empirical research suggested that the common usage of inappropriate names could produce a vicious circle in public communication and institutional notions further reinforce collective stigmatization, discrimination, and prejudice [45,72].

### Looking back and looking forward

Framed within the historical coevolution of scientific contexts, understanding the nosological continuity of diseases remains limit [63,77–80]. As a case in point, the pathological associations between *German measles* and common synonyms (e.g., *morbilli, rubeola, rubella, Rötheln*, etc.) are in the fog of confusion, although the debate has been going on over a century and a half earlier [7,81–85]. These diachronic discourses and lexical dynamics also remain unclear [4,7,86–89].

Nowadays, the Google Books Ngram Corpus (GBNC) is a unique linguistic landscape that benefits from centuries of development of rich grammatical and lexical resources, as well as its cultural context [27,90]. Arguably, the lexicographical and historiographical study promises to articulate the ins and outs of scientific narratives by leveraging the capacity of these rich metadata corpora over four centuries. As shown in **Figure 3**, many miscellaneous disease names (e.g., *morbilli, morbilli scarlatinosi, rötheln, feuermasern, scarlatina morbillosa, rubeola notha, rosalia idiopathica, bastard measles* or *scarlatina, hybrid measles* or *scarlatina*, etc.) have sunk back into merited oblivion in the ups and downs of epic history, whereas *German measles* was destined to become an antiquated and anachronistic usage, and *rubella* initiated a herald wave of dominant place after 1944.

**Figure 3.**
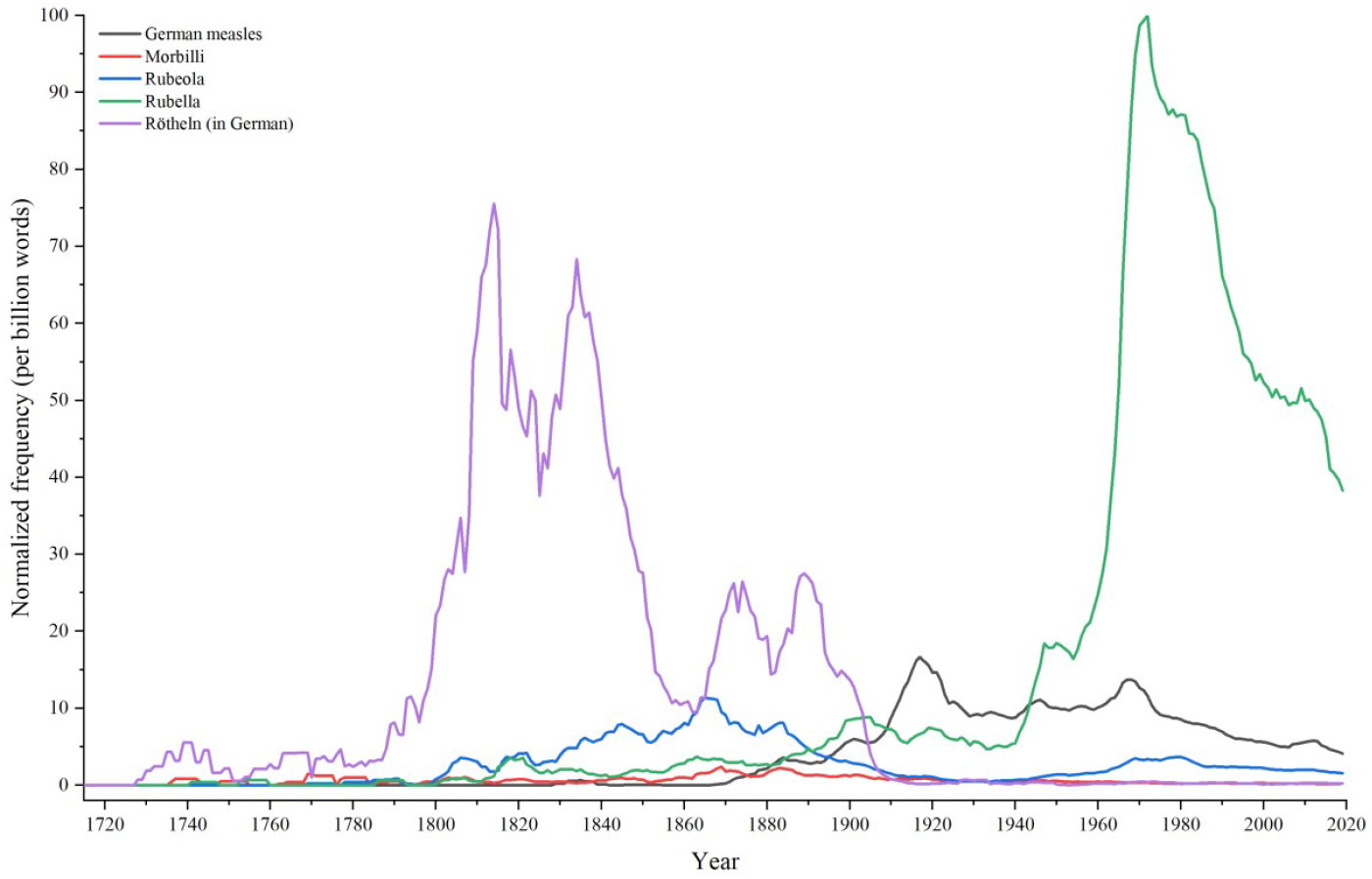
Historiographical study. Google Books Ngram Corpus (GBNC) facsimiles the diachronic discourse of *morbilli* (English corpus), *rubeola* (English corpus), *rubella* (English corpus), *Rötheln* (German corpus), and *German measles* (English corpus) from 1719 to 2019.

The nosology of *German measles* and similar diseases is still far from being generally recognized, as well as their pathological differentials [64,91]. *Measles* is an old English disease name that classical nosologists have vainly attempted to replace by such synonyms as *morbilli* and *rubeola* [92]. The English term *measles* was introduced by Dr. John of Gaddesden as an equivalent of the Latin term *morbilli* around the 14^th^ century [63,93,94]. But such designation was generally criticized for “a product of semantic and nosographic confusion.”[95] The term *rubeola* originally borrowed from the Latin word *Rubeus* (meaning *reddish*) in Avicenna of Bagdad’s writings, is thought to have been used for the first time as a translation of the term *measles* [94,96]. Indeed, the great majority of scientists recognize *German measles* to be an independent disease.

According to the OED Online, the earliest known references to *German measles* date back as far as 1856 (**Table 2**). Therefore, it is generally believed that the epidemic entity *German measles* was accepted growly after 1856 [7,97,98]. But this is not the case. The earliest usages could be stemmed back to about 1814 (**Table 3**).

**Table 2.**
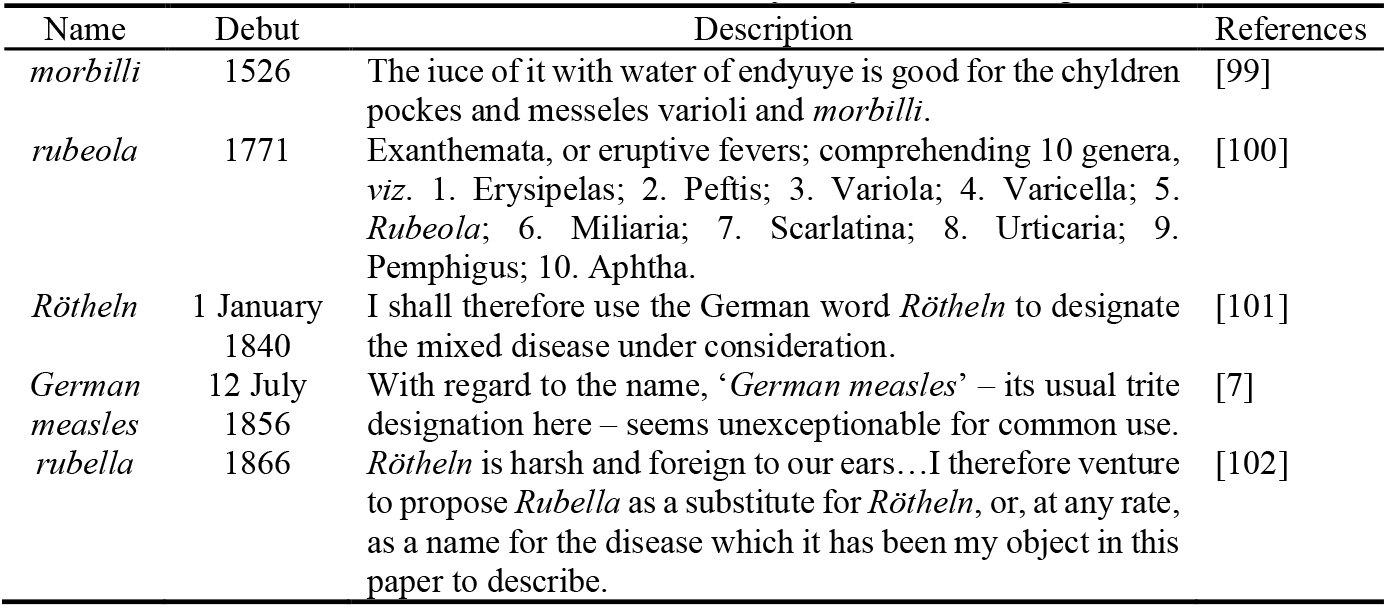
The debuts of *German measles* and its synonyms according to OED Online.

| Name | Debut | Description | References |
| --- | --- | --- | --- |
| *morbilli* | 1526 | The iuce of it with water of endyuye is good for the chyldren pockes and messeles varioli and *morbilli*. | [99] |
| *rubeola* | 1771 | Exanthemata, or eruptive fevers; comprehending 10 genera, viz. 1. Erysipelas; 2. Peftis; 3. Variola; 4. Varicella; 5. *Rubeola*; 6. Miliaria; 7. Scarlatina; 8. Urticaria; 9. Pemphigus; 10. Aphtha. | [100] |
| *Rötheln* | 1 January 1840 | I shall therefore use the German word *Rötheln* to designate the mixed disease under consideration. | [101] |
| *German measles* | 12 July 1856 | With regard to the name, ‘*German measles*’ – its usual trite designation here – seems unexceptionable for common use. | [7] |
| *rubella* | 1866 | *Rötheln* is harsh and foreign to our ears...I therefore venture to propose *Rubella* as a substitute for *Rötheln*, or, at any rate, as a name for the disease which it has been my object in this paper to describe. | [102] |

**Table 3.** Historiographical origins of *German measles* and common synonyms.

| Name | Debut | Credit | Evidence | References |
| --- | --- | --- | --- | --- |
| *rubeola* | 1768 | François Boissier de Sauvages de Lacroix (12 May 1706 – 19 February 1767) | Shortly before (1768), the two diseases had been separated by Sauvages in his Nosology, and he was the first to call measles “*rubeola*,” instead of “*morbilli*,” by which name it had always been known before. This new name, “*rubeola*,” was adopted by Cullen in his Nosology, published four years later (1772). | [89,103] |
| *Rötheln* | 1 January 1840 | Robert Paterson (1814 – 1889) | I fear that the adoption of the word *rubeola* for this disease would produce confusion in medical nomenclature. I shall therefore use the German word *Rötheln* to designate the mixed disease under consideration, in preference to that of *rubeola*, or the use of a new term. | [101,104] |
| *German measles* | 4 April 1814 | William George Maton (1774 – 1835) | On April 4, 1814, Dr. George Maton ... This first identification of *German measles* as a discrete illness was published one year later, an interval from presentation to publication not dissimilar to that in modern experience. | [105–107] |
| *rubella* | 1740 | Friedrich Hoffmann (1660 – 1742) | Friedrich Hoffmann (1660-1742), ..., Notable among his many clinical descriptions are those of *rubella* (called “German” measles as a consequence of his description,) chlorosis, and the diseases of the pancreas and liver. | [108,109] |

The term *German Measles* was established as a separate disease in 1814 and officially recognized by the International Congress of Medicine in 1881. The known clinical description came from German physicians Friedrich Hoffmann in 1740, De Bergen in 1752, and Orlow in 1758, respectively [97,108,109]. Before 1768, for more learned occasions, *Rötheln* and *morbilli* seem more decidedly to mark a distinct disease, than any other yet proposed [7,89]. French physician Sauvages de Lacroix, who established the first methodical nosology for disease classification in 1763 [57,110], first applied the term *rubeola* to what had been previously termed *morbilli* in 1768 [89]. And while almost immediately after him, the German physicians, Selle, Orlow, and Ziegler, clearly laid down the distinctive marks between *rubeola* and *morbilli*. On April 4, 1814, Dr. George de Maton read a paper entitled “*Some Account of a Rash Liable to be Mistaken for Scarlatina*” at the Royal College of Physicians in London [105–107], which results in the names *rubella* or *German measles* as a substitute for *Rötheln* [7,86]. Then, the epidemic term *German measles* was accepted gradually as a synonym of *rubella. German measles, Rötheln* or *rubeola* per se, was officially ratified as a distinct disease at the 7^th^ International Medical Congress, London, August 2 to 9, 1881 [88,111–118]. A quarter-century later, the term *German Measles* has ultimately become common usage.

*Rubella* has been “discovered – and named – multiple times” in the past centuries [119]. In modern literature, *rubella* has become a *de facto* synonym for *German Measles* after 1944 [7–16]. In 1740, the English name *rubella* is derived from Latin *rubellus* reddish, and the clinical description of *rubella* was first described by Friedrich Hoffmann, the author of *Fundamenta Medicinae* [108,109]. Then, *rubella* was considered by Dr. Maton to be a mere variant of measles or scarlet fever in 1814 [105,106,120]. Half a century later, English surgeon Henry Veale suggested the need to name the discrete disease, and formally proposed the name *rubella* as a substitute for *Rötheln* in 1866 [97]. As a major human infectious disease, *rubella* must have emerged only in the past 11,000 years for which some close relatives may still exist among animals [4,64]. Indeed, consistent with the historiographical results (**Figure 3**), *rubella* had been considered of “minor importance among the common communicable diseases” until 1940 [121]. Following the *rubella* epidemic of 1940, the occurrence of congenital rubella syndrome (CRS) was first recognized by Norman McAlister Gregg in 1941 [122,123]. As of 2018, 81 countries were verified as having eliminated *rubella* via routine vaccination, and even today *rubella* remains endemic in other countries [124].

### A heuristic roadmap of exclusive substitute

An exclusive substitution for an anachronistic usage could be a pre-existed synonym, a blend word, or a neologism. However, we should curate an exclusive substitute following the ethical principles (**Figure 2**). Relatedly, as a heuristic case, we hash out the inappropriate name like *German Measles* to quell confusion and avoid stigma. Here, we demonstrate an illustrational approach to determine an exclusive substitution for *German Measles* without ambiguity.

First, the similarity coefficient between words is determined by deep learning models, and finally screen out an exclusive substitute for *German Measles* according to the semantic similarity scores of word embeddings. In **Figure 4**, the input example is first constructed by summing the corresponding token, segment, and position embeddings, and then word embeddings go through the BERT base model to obtain the vector representations with semantic information. As for the bigram like *German measles*, we averaged the individual word vector to get the final word vector. By quantifying the cosine similarity scores between the word vectors (**Figure 5**), it turns out that the term *rotheln* is substantial equivalence to *rötheln* with the highest semantic fidelity, and the results of the BioBERT model and the PubMedBERT model shed light on each other. Most notably, as a model in the medical field, the PubMedBERT model maps out that *Rubella* should be the exclusive substitution for *German measles* with the highest semantic similarity.

**Figure 4.**
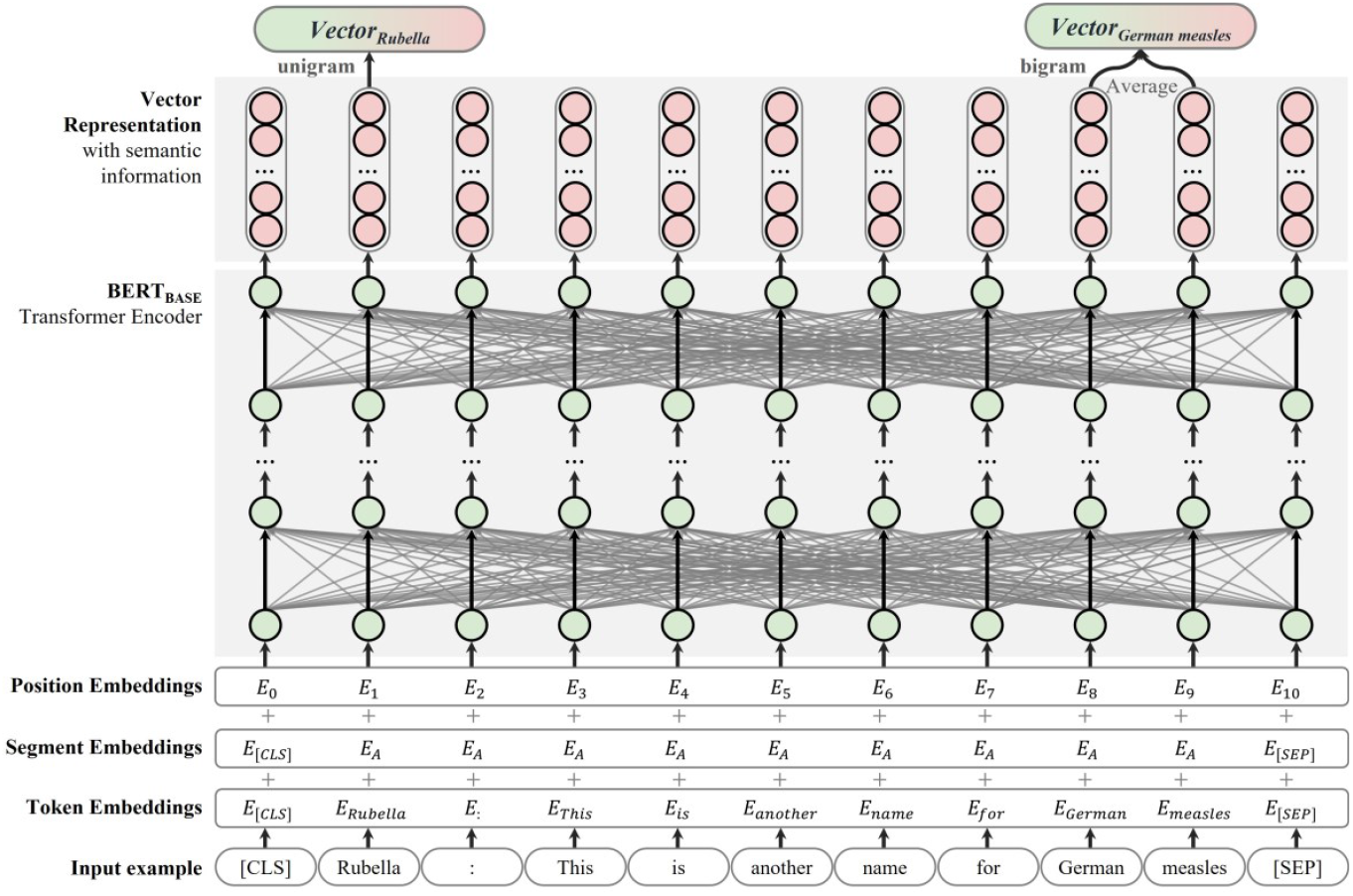
Illustrational architecture of the BERT and PubMedBERT models.

**Figure 5.**
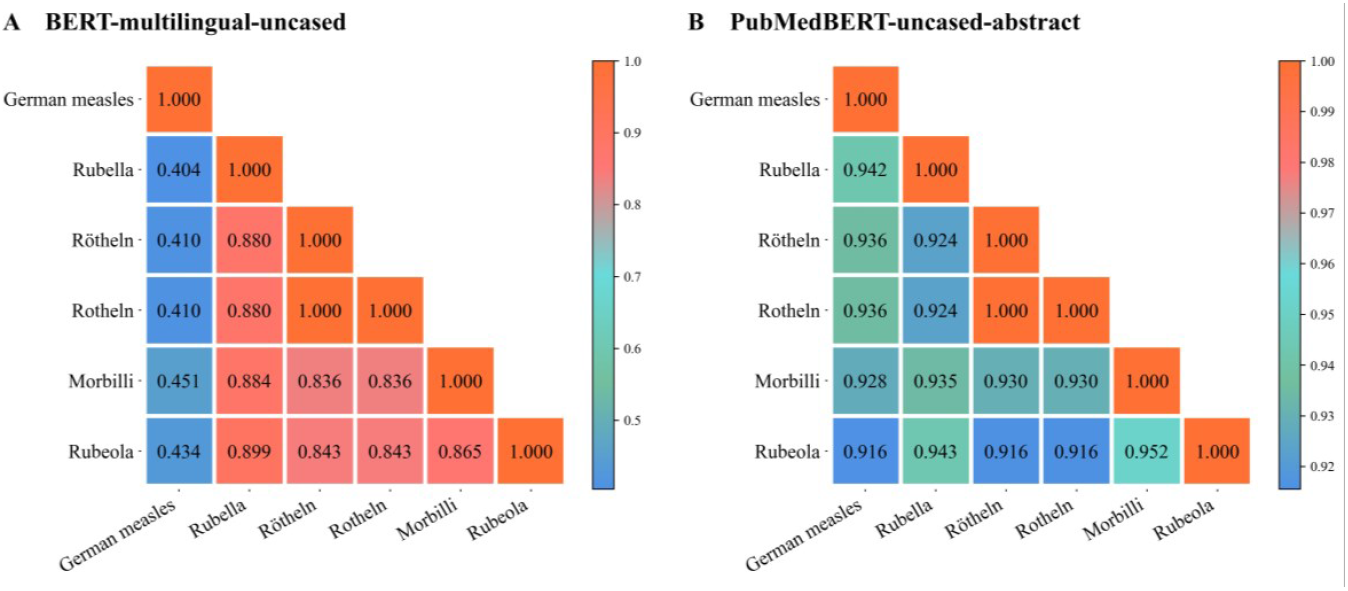
Heatmaps of semantic similarity scores. A, the BERT model and B, the PubMedBERT model. The higher scores in the heatmaps, the higher the semantic similarity between two synonyms.

Second, some case studies are given using the same function with the PubMedBERT model for *post-hoc* explanations (**Table 4**). According to syntactical function, in case #1, the qualifier ‘this’ refers to *rubella*, so the synonymous sentence of the original sentence was that *rubella* is another name for *German measles*. Semantically, *rubella* has an equivalence relationship with *German measles*, with a very high semantic relevance (0.954). In case #2, *rötheln* has a dependent relationship with *German measles* rather than an equivalence relationship. In cases #3 and #4, *morbilli* and *rubeola* tend to the appositions of *measles* rather than those of *German measles*. To sum up, the case studies further emphasize the high semantic similarity between *rubella* and *German measles*.

**Table 4.**
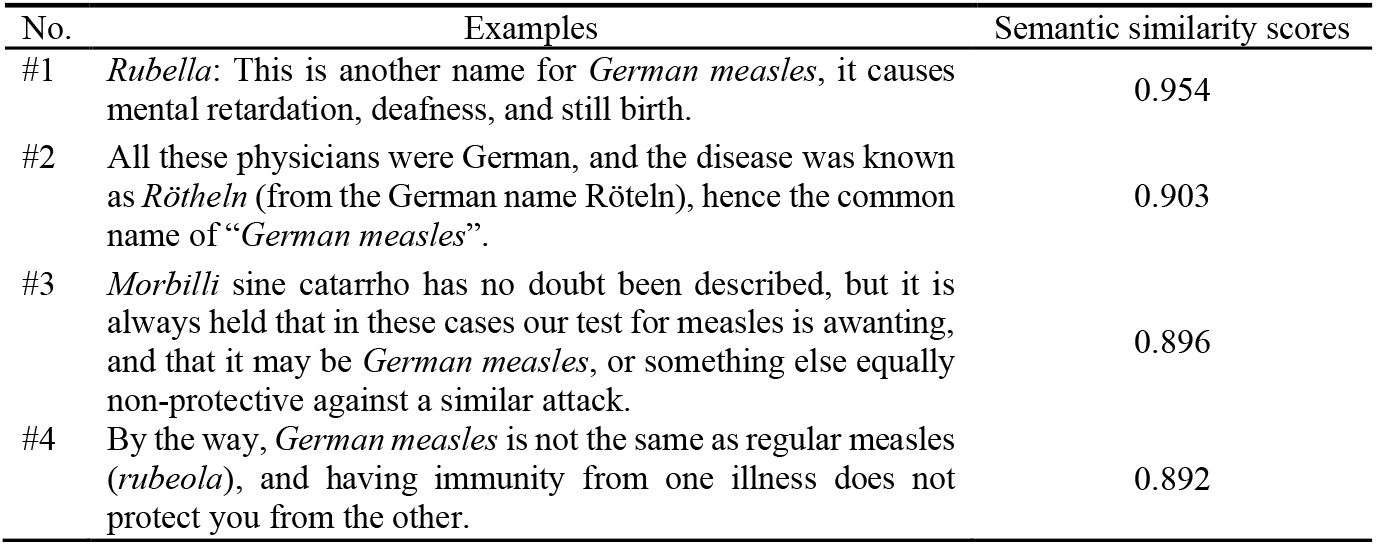
Some case studies are retrieved using the same function with the PubMedBERT model.

Third, the purpose of the semantic drift experiments is to examine how and to what extent the meanings of *German measles, morbilli, rubeola, rubella*, and *rötheln* have changed over time. In the meantime, estimate the semantics of words from a particular period through historical synonyms. In **Figure 6**, we projected the latent semantic drifts of *German measles, morbilli, rubeola, rubella*, and *rötheln* from their debuts to 2020 (**Table 2** and **Table 3**). To highlight the cases of semantic drift, the five keywords (in colors) were shown with their historical synonyms (in gray). The closer two words are to each other, the more semantically similar they are. For example, the term *Rubella* was closely associated with the German term *masern* in 1740, and the dominant meaning of *Rubella* was more semantically similar to the words *fowlpox* and *morbillivirus* in 2020. As for the term *German measles*, its meaning was closer to *pustula, flowered measles*, and *strawberry measles* when it debuted. However, the semantics of *German measles* is closer to those of *paramyxoviridae* and *three day measles* today. *Rubeola* changed its dominant meaning from skin disease *scabies* to *röteln*, and this change approximately took place between 1768 and 2020. Comparatively, *Rötheln* was more often associated with the word *variola* rather than *anthrax*, while *Morbilli* was referred to *pediculosis* or *rash* rather than *rosacea* or *red sandwort*. Therefore, it is found that five keywords have different degrees of semantic drift over time. Coupled with the previous results (**Figure 3**), *rubella* is a high-pragmatic-frequency synonym of *German Measles* in recent literature and tends to survive in due course.

**Figure 6.**
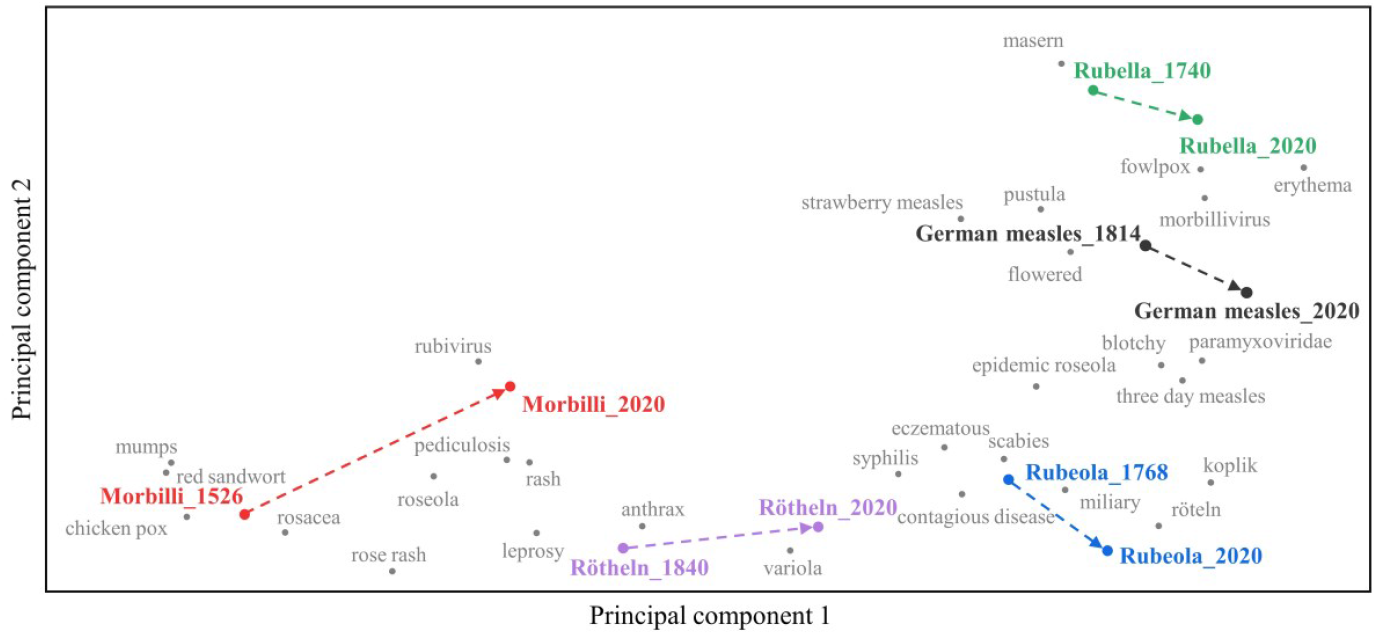
The historical reconceptualization process of *German measles* and common synonyms.

In short, our results strongly suggest that *rubella* is a geography-free, high-pragmatic-frequency, and high-semantic-homology synonym of *German Measles*. In theory, *rubella* is a pithy substitute for efficient communication according to Zipf’s principle of least effort, comparing with the blend-word *German Measles* [125]. Additionally, it rolls much easier off the tongue than *German Measles* in daily communication. In retrospect, some pioneers advocated the discarding of the offensive name *German Measles* before [91,126,127], as the remarks, “it [*rubella*] is perhaps the best that has been used”[91] and “a better name for which [*German Measles*] is *rubella*.”[127] Such foresight is also consistent with our experimental results. Therefore, it should be an optimal substitute to fill in the niche of *German Measles* in practice.

## Discussion

### Conclusion

On the anniversary of the best practices of new human infectious diseases announced by WHO, we take an open mind to appreciate modest introspections and rededications to celebrate the big moment. The present naming rules could not offer a one-size-fits-all corrective mechanism, especially for the pre-existed names. In the scientific sphere, much remains to be learned about the ins and outs of naming practices. Here, we examine what we need to know and what we need to eliminate the label paradox in due course.

Concretely, our infodemiological study first shows that long-standing but flawed names of human diseases are still going viral in both the scientific community and news outlets at the cost of social impacts, whatever their seemingly harmless origins. Following the best practices of WHO, curated names of human diseases should be scientifically pithy and socially acceptable, with the faith of minimizing marginal impacts on nations, economies, and people.

Our lexicographical and historiographical study could articulate the twists and turns of naming human diseases over several centuries, penetrate to the essence of nosology, and finally bridge the gaps of contemporary considerations. Heuristic introspection would help us to determine pithy names instead of offensive counterparts. Arguably, as an exemplificative case, it is reasonable that *rubella* should become an exclusive usage substitute for *German Measles* with the same clinical manifestations and equivalent semantics. Thus, our proposed principles and approaches are expected to provide hallmark remedial mechanisms for current nosology-based standards and reframe far-reaching discussions on the nomenclature of human diseases.

## Acknowledgements

We hereby desire to express indebtedness to anonymous reviewers for their valuable and constructive comments. This study was partially supported by the National Natural Science Foundation of China under Grant U1936208, the Zhejiang Provincial Natural Science Foundation of China under Grant LZ21F020004, and the Open Research Project of the State Key Laboratory of Media Convergence and Communication, Communication University of China under Grant SKLMCC2021KF015.

## Data availability

The synthetic data generated in this study and custom code supporting this study are available at GitHub (https://github.com/YaChen8/Naming_human_disease).

## Abbreviations

BERT: Bidirectional Encoder Representations from Transformers
COVID-19: *Coronavirus* Disease 2019
GBNC: Google Books Ngram Corpus
GDELT: Global Data on Events, Location and Tone
ICD: International Classification of Diseases
ICD-11: Eleventh revision of the International Classification of Diseases
MERS: Middle Eastern Respiratory Syndrome
OED Online: Oxford English Dictionary Online
PCA: Principal Component Analysis
PPMI: Positive Pointwise Mutual Information
SARS: Severe Acute Respiratory Syndrome
SVD: Singular Value Decomposition
WHA: World Health Assembly
WHO: World Health Organization

## Multimedia Appendix 1 PRISMA-based flowchart

In this study, we orchestrate rich metadata available to unveil the scientific paradigms via four pertinent experiments, following the Preferred Reporting Items for Systematic Reviews and Meta-Analyses (PRISMA) guidelines (PRISMA 2015; Zeraatkar and Ahmadi 2018; Jobin et al. 2019; Page et al. 2021).

**Figure A1.**
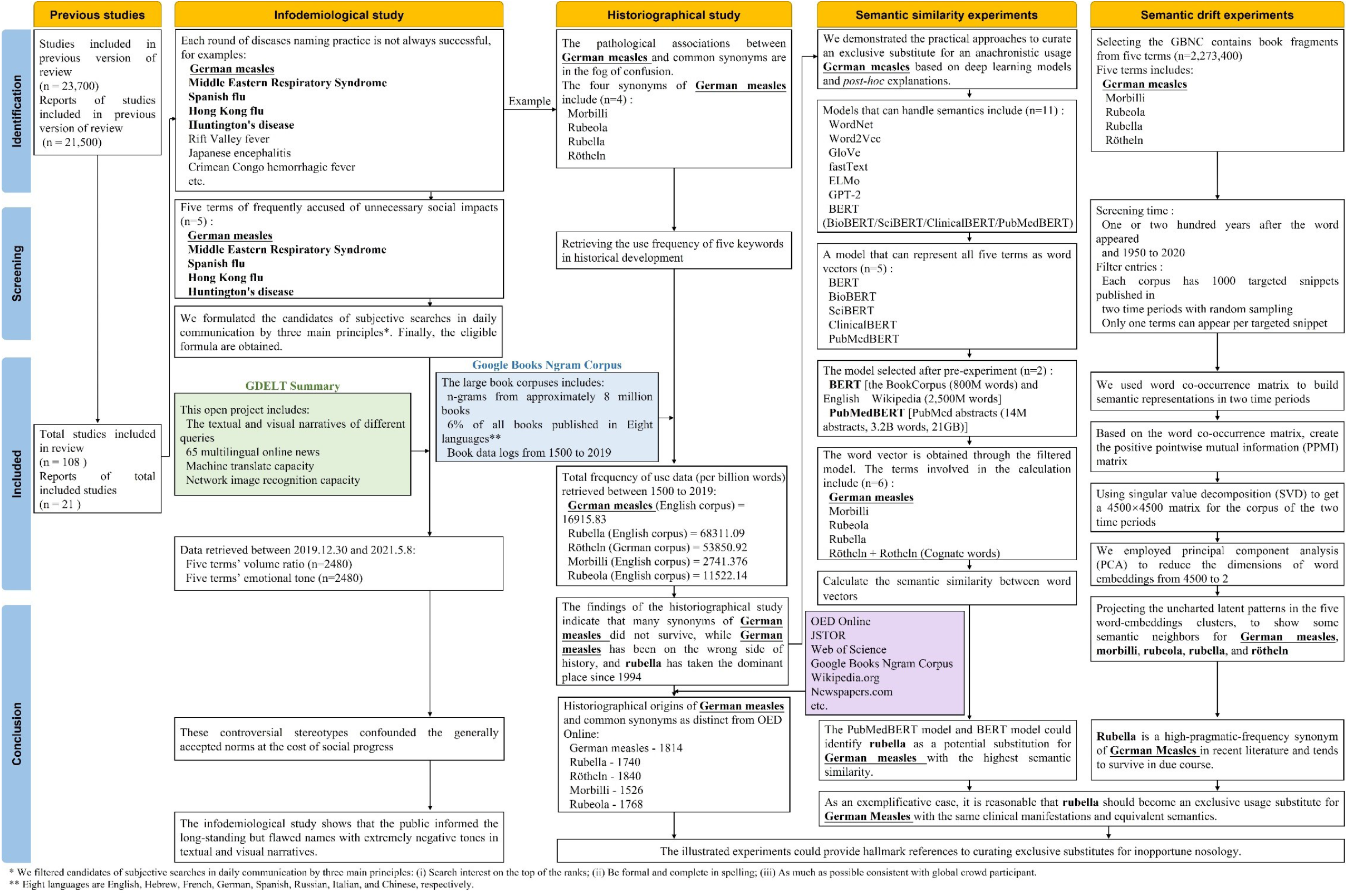
PRISMA-based flowchart.

## Multimedia Appendix 2 Code scheme

A defined itemized code scheme of crowd behavior in daily communication is paramount for our understanding of the unbiased and comprehensive archive of global online news. Firstly, the initial search candidates included *German measles, Middle Eastern Respiratory Syndrome, Spanish flu, Hong Kong flu*, and *Huntington’s disease*. They have been accused of unnecessary social impacts before [1–10]. Then, we formulated the candidates of subjective searches in daily communication by three main principles: (i) Search interest on the top of the ranks; (ii) Be formal and complete in spelling; (iii) As much as possible consistent with global crowd participant [11]. Finally, the eligible search formulas meeting the inclusion criteria are as following:

1. German measles: *(“German measles” OR “German Measles”*) *AND PublicationDate>=12/30/2019 AND PublicationDate<=5/8/2021*
2. Middle Eastern Respiratory Syndrome: *(“Middle Eastern respiratory syndrome” OR “Middle Eastern Respiratory Syndrome”*) *AND PublicationDate>=12/30/2019 AND PublicationDate<=5/8/2021*
3. Spanish flu: *(“Spanish flu” OR “Spanish Flu” OR “Spanish influenza” OR “Spanish Influenza”*) *AND PublicationDate>=12/30/2019 AND PublicationDate<=5/8/2021*
4. Hong Kong flu: *(“Hong Kong flu” OR “Hong Kong Flu” OR “Hong Kong influenza” OR “Hong Kong Influenza”*) *AND PublicationDate>=12/30/2019 AND PublicationDate<=5/8/2021*
5. Huntington’s disease: *(“Huntington’s disease” OR “Huntington’s chorea” OR “huntington’s disease” OR “huntington’s chorea” OR “Huntington Disease” OR “Huntington disease” OR “Huntington chorea” OR “huntington disease” OR “huntington chorea”*) *AND PublicationDate>=12/30/2019 AND PublicationDate<=5/8/2021*

By leveraging the capacity of GDELT’s machine translate and neural network image recognition, the unbiased news portfolio in 65 languages provides a unique lens into global online news coverage from December 30, 2019 to May 8, 2021.

## References

1. WHO. World Health Organization Best Practices for the Naming of New Human Infectious Diseases [Internet]. 2015. p. 1–3. Available from: https://www.who.int/topics/infectious_diseases/naming-new-diseases/en/

2. Fukuda K, Wang R, Vallat B. Naming diseases: First do no harm. Science [Internet] 2015 May 8;348(6235) :643–643. [doi: 10.1126/science.348.6235.643]

3. Chandrashekhar V. The burden of stigma. Science [Internet] 2020 Sep 18;369(6510) :1419–1423. [doi: 10.1126/science.369.6510.1419]

4. Bennett AJ, Paskey AC, Ebinger A, Pfaff F, Priemer G, Höper D, Breithaupt A, Heuser E, Ulrich RG, Kuhn JH, Bishop-Lilly KA, Beer M, Goldberg TL. Relatives of rubella virus in diverse mammals. Nature [Internet] 2020 Oct 15;586(7829) :424–428. [doi: 10.1038/s41586-020-2812-9]

5. Gibbons A. Newly found viruses suggest rubella originated in animals. Science [Internet] 2020 Oct 9;370(6513) :157–157. [doi: 10.1126/science.370.6513.157]

6. Gibbons A. Newly discovered viruses suggest ‘German measles’ jumped from animals to humans. Science [Internet] 2020 Oct 7; [doi: 10.1126/science.abf1520]

7. Philo Scotus. The Rubeola Epidemic. Lancet [Internet] 1856 Jul;68(1715) :57. [doi: 10.1016/S0140-6736(02)76420-9]

8. Wesselhoeft C. Rubella (German Measles). N Engl J Med [Internet] 1947 Jun 19;236(25) :943–950. [doi: 10.1056/NEJM194706192362506]

9. Wesselhoeft C. Rubella (German Measles). N Engl J Med [Internet] 1947 Jun 26;236(26) :978–988. [doi: 10.1056/NEJM194706262362605]

10. Wesselhoeft C. Rubella (German Measles) and Congenital Deformities. N Engl J Med [Internet] 1949 Feb 17;240(7) :258–261. [doi: 10.1056/NEJM194902172400706]

11. Dudgeon JA. Immunization against Rubella. Nature [Internet] 1969 Aug;223(5207) :674–676. [doi: 10.1038/223674a0]

12. Eichhorn MM. Rubella: Will Vaccination Prevent Birth Defects? Science [Internet] 1971 Aug 20;173(3998) :710–711. [doi: 10.1126/science.173.3998.710-a]

13. Eichhorn MM. Rubella: Will Vaccination Prevent Birth Defects? Science [Internet] 1971 Aug 20;173(3998) :710–711. [doi: 10.1126/science.173.3998.710-b]

14. Donald MW, Goff WR. Attention-Related Increases in Cortical Responsivity Dissociated from the Contingent Negative Variation. Science [Internet] 1971 Jun 11;172(3988) :1163–1166. [doi: 10.1126/science.172.3988.1163]

15. Maugh TH. Diabetes: Epidemiology Suggests a Viral Connection. Science [Internet] 1975 Apr 25;188(4186) :347–351. [doi: 10.1126/science.188.4186.347]

16. Gordis L, Gold E. Privacy, confidentiality, and the use of medical records in research. Science [Internet] 1980 Jan 11;207(4427) :153–156. [doi: 10.1126/science.7350648]

17. Jardetzky TS, Lamb RA. A class act. Nature [Internet] 2004 Jan;427(6972) :307–308. [doi: 10.1038/427307a]

18. Wadman M. The physician whose 1964 vaccine beat back rubella is working to defeat the new coronavirus. Science [Internet] 2020 Mar 21; [doi: 10.1126/science.abb8290]

19. Shimizu K. 2019-nCoV, fake news, and racism. Lancet [Internet] 2020 Feb;395(10225) :685–686. [doi: 10.1016/S0140-6736(20)30357-3]

20. The Lancet. COVID-19: fighting panic with information. Lancet [Internet] 2020 Feb;395(10224) :537. [doi: 10.1016/S0140-6736(20)30379-2]

21. Lazer D, Pentland A, Adamic L, Aral S, Barabasi A-L, Brewer D, Christakis N, Contractor N, Fowler J, Gutmann M, Jebara T, King G, Macy M, Roy D, Van Alstyne M. Computational Social Science. Science [Internet] 2009 Feb 6;323(5915) :721–723. [doi: 10.1126/science.1167742]

22. Lazer DMJ, Pentland A, Watts DJ, Aral S, Athey S, Contractor N, Freelon D, Gonzalez-Bailon S, King G, Margetts H, Nelson A, Salganik MJ, Strohmaier M, Vespignani A, Wagner C. Computational social science: Obstacles and opportunities. Science [Internet] 2020 Aug 28;369(6507) :1060–1062. [doi: 10.1126/science.aaz8170]

23. Page MJ, Moher D, Bossuyt PM, Boutron I, Hoffmann TC, Mulrow CD, Shamseer L, Tetzlaff JM, Akl EA, Brennan SE, Chou R, Glanville J, Grimshaw JM, Hróbjartsson A, Lalu MM, Li T, Loder EW, Mayo-Wilson E, McDonald S, McGuinness LA, Stewart LA, Thomas J, Tricco AC, Welch VA, Whiting P, McKenzie JE. PRISMA 2020 explanation and elaboration: updated guidance and exemplars for reporting systematic reviews. BMJ [Internet] 2021 Mar 29;n160. [doi: 10.1136/bmj.n160]

24. Hu Z, Yang Z, Li Q, Zhang A. The COVID-19 Infodemic: Infodemiology Study Analyzing Stigmatizing Search Terms. J Med Internet Res [Internet] 2020 Nov 16;22(11) :e22639. PMID:33156807

25. Leetaru KH, Schrodt PA. A 30-Year Georeferenced Global Event Database: The Global Database of Events, Language, and Tone (GDELT). 54th Annu Conv Int Stud Assoc [Internet] San Francisco, California, USA; 2013. p. 1–51. Available from: http://data.gdeltproject.org/documentation/ISA.2013.GDELT.pdf

26. Wang W, Kennedy R, Lazer D, Ramakrishnan N. Growing pains for global monitoring of societal events. Science [Internet] 2016 Sep 30;353(6307) :1502–1503. [doi: 10.1126/science.aaf6758]

27. Michel J-B, Shen YK, Aiden AP, Veres A, Gray MK, Pickett JP, Hoiberg D, Clancy D, Norvig P, Orwant J, Pinker S, Nowak MA, Aiden EL. Quantitative Analysis of Culture Using Millions of Digitized Books. Science [Internet] 2011 Jan 14;331(6014) :176–182. [doi: 10.1126/science.1199644]

28. Devlin J, Chang M-W, Lee K, Toutanova K. BERT: Pre-training of Deep Bidirectional Transformers for Language Understanding. Proc 2019 Conf North Am [Internet] Stroudsburg, PA, USA: Association for Computational Linguistics; 2019. p. 4171–4186. [doi: 10.18653/v1/N19-1423]

29. Gu Y, Tinn R, Cheng H, Lucas M, Usuyama N, Liu X, Naumann T, Gao J, Poon H. Domain-Specific Language Model Pretraining for Biomedical Natural Language Processing. 2020 Jul 30; Available from: http://arxiv.org/abs/2007.15779

30. Li Y, Engelthaler T, Siew CSQ, Hills TT. The Macroscope: A tool for examining the historical structure of language. Behav Res Methods [Internet] 2019 Aug 11;51(4) :1864–1877. [doi: 10.3758/s13428-018-1177-6]

31. Li Y, Hills T, Hertwig R. A brief history of risk. Cognition [Internet] 2020 Oct;203:104344. [doi: 10.1016/j.cognition.2020.104344]

32. Kupferschmidt K. Rules of the name. Science [Internet] 2015 May 15;348(6236) :745–745. [doi: 10.1126/science.348.6236.745]

33. Kupferschmidt K. Discovered a disease? WHO has new rules for avoiding offensive names. Science [Internet] 2015 May 11; [doi: 10.1126/science.aac4575]

34. Enserink M. Amid Heightened Concerns, New Name for Novel Coronavirus Emerges. Science [Internet] 2013 May 10;340(6133) :673–673. PMID:23661733

35. Dowell SF, Bresee JS. Pandemic lessons from Iceland. Proc Natl Acad Sci [Internet] 2008 Jan 29;105(4) :1109–1110. [doi: 10.1073/pnas.0711535105]

36. Hoppe T. “Spanish Flu”: When Infectious Disease Names Blur Origins and Stigmatize Those Infected. Am J Public Health [Internet] 2018 Nov;108(11) :1462–1464. [doi: 10.2105/AJPH.2018.304645]

37. Anonymous. Have we had Hong Kong flu before? Lancet [Internet] 1969 May;293(7601) :929–930. [doi: 10.1016/S0140-6736(69)92556-2]

38. Peckham R. Viral surveillance and the 1968 Hong Kong flu pandemic. J Glob Hist [Internet] 2020 Nov 6;15(3) :444–458. [doi: 10.1017/S1740022820000224]

39. Chang, BS C, Salerno, MSc A, Hsu, MD, MPH EB. Perspectives on xenophobia during epidemics and implications for emergency management. J Emerg Manag [Internet] 2020 Dec 10;18(7) :23–29. [doi: 10.5055/jem.2020.0521]

40. Wexler A. Stigma, history, and Huntington’s disease. Lancet [Internet] 2010 Jul;376(9734) :18–19. [doi: 10.1016/S0140-6736(10)60957-9]

41. Rawlins M. Huntington’s disease out of the closet? Lancet [Internet] 2010 Oct;376(9750) :1372–1373. [doi: 10.1016/S0140-6736(10)60974-9]

42. Spinney L. Uncovering the true prevalence of Huntington’s disease. Lancet Neurol [Internet] 2010 Aug;9(8) :760–761. [doi: 10.1016/S1474-4422(10)70160-5]

43. Editorial. Dispelling the stigma of Huntington’s disease. Lancet Neurol [Internet] 2010 Aug;9(8) :751. [doi: 10.1016/S1474-4422(10)70170-8]

44. Hyman SE. The Unconscionable Gap Between What We Know and What We Do. Sci Transl Med [Internet] 2014 Sep 10;6(253) :1–4. [doi: 10.1126/scitranslmed.3010312]

45. Gollwitzer A, Marshall J, Wang Y, Bargh JA. Relating pattern deviancy aversion to stigma and prejudice. Nat Hum Behav [Internet] 2017 Dec 27;1(12) :920–927. [doi: 10.1038/s41562-017-0243-x]

46. Soroka S, Fournier P, Nir L. Cross-national evidence of a negativity bias in psychophysiological reactions to news. Proc Natl Acad Sci [Internet] 2019 Sep 17;116(38) :18888–18892. [doi: 10.1073/pnas.1908369116]

47. Pescosolido BA, Martin JK. The Stigma Complex. Annu Rev Sociol [Internet] 2015 Aug 14;41(1) :87–116. [doi: 10.1146/annurev-soc-071312-145702]

48. WHO. WHO issues best practices for naming new human infectious diseases. 2015; Available from: https://www.who.int/news/item/08-05-2015-who-issues-best-practices-for-naming-new-human-infectious-diseases

49. WHO. A guide to preventing and addressing social stigma [Internet]. Available from: https://www.who.int/docs/default-source/coronaviruse/covid19-stigma-guide.pdf

50. Editorial. Stop the coronavirus stigma now. Nature [Internet] 2020 Apr 9;580(7802) :165–165. [doi: 10.1038/d41586-020-01009-0]

51. London AJ, Kimmelman J. Against pandemic research exceptionalism. Science [Internet] 2020 May 1;368(6490) :476–477. [doi: 10.1126/science.abc1731]

52. Olson DR, Simonsen L, Edelson PJ, Morse SS. Epidemiological evidence of an early wave of the 1918 influenza pandemic in New York City. Proc Natl Acad Sci [Internet] 2005 Aug 2;102(31) :11059–11063. [doi: 10.1073/pnas.0408290102]

53. Editorial. Calling it what it is. Nat Genet [Internet] 2020 Apr 3;52(4) :355–355. [doi: 10.1038/s41588-020-0617-2]

54. Fischer LS, Mansergh G, Lynch J, Santibanez S. Addressing Disease-Related Stigma During Infectious Disease Outbreaks. Disaster Med Public Health Prep [Internet] 2019 Dec 3;13(5–6): 989–994. [doi: 10.1017/dmp.2018.157]

55. WHO. Ebola haemorrhagic fever in Zaire, 1976. Bull World Health Organ [Internet] 1978 Sep 12;56(2) :271–93. PMID:307456

56. WHO. Reference Guide on the Content Model of the ICD-11 alpha [Internet]. 2011. p. 59. Available from: https://www.who.int/classifications/icd/revision/Content_Model_Reference_Guide.January_2011.pdf

57. Kveim Lie A, Greene JA. From Ariadne’s Thread to the Labyrinth Itself — Nosology and the Infrastructure of Modern Medicine. Malina D, editor. N Engl J Med [Internet] 2020 Mar 26;382(13) :1273–1277. [doi: 10.1056/NEJMms1913140]

58. Fortunato S, Bergstrom CT, Börner K, Evans JA, Helbing D, Milojević S, Petersen AM, Radicchi F, Sinatra R, Uzzi B, Vespignani A, Waltman L, Wang D, Barabási A-L. Science of science. Science [Internet] 2018 Mar 2;359(6379) :eaao0185. [doi: 10.1126/science.aao0185]

59. Enserink M. War Stories. Science [Internet] 2013 Mar 15;339(6125) :1264–1268. [doi: 10.1126/science.339.6125.1264]

60. Honigsbaum M. Jean-Jacques Muyembe-Tamfum: Africa’s veteran Ebola hunter. Lancet [Internet] 2015 Jun;385(9986) :2455. [doi: 10.1016/S0140-6736(15)61128-X]

61. Jean-Jacques Muyembe Tamfum: a life’s work on Ebola. Bull World Health Organ [Internet] 2018 Dec 1;96(12) :804–805. [doi: 10.2471/BLT.18.031218]

62. Kuhn JH, Adachi T, Adhikari NKJ, Arribas JR, Bah IE, Bausch DG, Bhadelia N, Borchert M, Brantsæter AB, Brett-Major DM, Burgess TH, Chertow DS, Chute CG, Cieslak TJ, Colebunders R, Crozier I, Davey RT, de Clerck H, Delgado R, Evans L, Fallah M, Fischer WA, Fletcher TE, Fowler RA, Grünewald T, Hall A, Hewlett A, Hoepelman AIM, Houlihan CF, Ippolito G, Jacob ST, Jacobs M, Jakob R, Jacquerioz FA, Kaiser L, Kalil AC, Kamara RF, Kapetshi J, Klenk H-D, Kobinger G, Kortepeter MG, Kraft CS, Kratz T, Bosa HSK, Lado M, Lamontagne F, Lane HC, Lobel L, Lutwama J, Lyon GM, Massaquoi MBF, Massaquoi TA, Mehta AK, Makuma VM, Murthy S, Musoke TS, Muyembe-Tamfum J-J, Nakyeyune P, Nanclares C, Nanyunja M, Nsio-Mbeta J, O’Dempsey T, Pawęska JT, Peters CJ, Piot P, Rapp C, Renaud B, Ribner B, Sabeti PC, Schieffelin JS, Slenczka W, Soka MJ, Sprecher A, Strong J, Swanepoel R, Uyeki TM, van Herp M, Vetter P, Wohl DA, Wolf T, Wolz A, Wurie AH, Yoti Z. New filovirus disease classification and nomenclature. Nat Rev Microbiol [Internet] 2019 May 29;17(5) :261–263. [doi: 10.1038/s41579-019-0187-4]

63. Sykes W. On the origin and history of some disease-names. Lancet [Internet] 1896 Apr;147(3789) :1007–1010. [doi: 10.1016/S0140-6736(01)39505-3]

64. Wolfe ND, Dunavan CP, Diamond J. Origins of major human infectious diseases. Nature [Internet] 2007 May;447(7142) :279–283. [doi: 10.1038/nature05775]

65. Trisnawati I, Budiono E, Sumardi, Setiadi A. Traumatic Inhalation due to Merapi Volcanic Ash. Acta Med Indones [Internet] 2015 Jul;47(3) :238–43. PMID:26586390

66. Burhan E, Mukminin U. A systematic review of respiratory infection due to air pollution during natural disasters. Med J Indones [Internet] 2020 Mar 20;29(1) :11–8. [doi: 10.13181/mji.oa.204390]

67. Kim Y. Renaming the term schizophrenia in Japan. Lancet [Internet] 2002 Sep;360(9336) :879. [doi: 10.1016/S0140-6736(02)09987-7]

68. Maruta T, Volpe U, Gaebel W, Matsumoto C, Iimori M. Should schizophrenia still be named so? Schizophr Res [Internet] 2014 Jan;152(1) :305–306. [doi: 10.1016/j.schres.2013.11.005]

69. Sartorius N. Stigma: what can psychiatrists do about it? Lancet [Internet] 1998 Sep;352(9133) :1058–1059. [doi: 10.1016/S0140-6736(98)08008-8]

70. Park J-H, Choi Y-M, Kim B, Lee D-W, Gim M-S. Use of the Terms “Schizophrenia” and “Schizophrenic” in the South Korean News Media: A Content Analysis of Newspapers and News Programs in the Last 10 Years. Psychiatry Investig [Internet] 2012;9(1) :17. [doi: 10.4306/pi.2012.9.1.17]

71. George B, Klijn A. A modern name for schizophrenia (PSS) would diminish self-stigma. Psychol Med [Internet] 2013 Jul 17;43(7) :1555–1557. [doi: 10.1017/S0033291713000895]

72. Porter R. Can the stigma of mental illness be changed? Lancet [Internet] 1998 Sep;352(9133) :1049–1050. [doi: 10.1016/S0140-6736(98)07155-4]

73. Corrigan PW. Erasing Stigma Is Much More Than Changing Words. Psychiatr Serv [Internet] 2014 Oct;65(10) :1263–1264. [doi: 10.1176/appi.ps.201400113]

74. Koike S, Yamaguchi S, Ojio Y, Ohta K, Ando S. Effect of Name Change of Schizophrenia on Mass Media Between 1985 and 2013 in Japan: A Text Data Mining Analysis. Schizophr Bull [Internet] 2016 May;42(3) :552–559. [doi: 10.1093/schbul/sbv159]

75. Corrigan PW. Beware the Word Police. Psychiatr Serv [Internet] 2019 Mar;70(3) :234–236. [doi: 10.1176/appi.ps.201800369]

76. Editorial. Embrace the WHO’s new naming system for coronavirus variants. Nature [Internet] 2021 Jun 10;594(7862) :149–149. [doi: 10.1038/d41586-021-01508-8]

77. Anonymous. The nomenclature of diseases: I. BMJ 1868;2(403) :316–318.

78. Anonymous. The nomenclature of diseases:II. BMJ 1869;2(406) :396–397.

79. Anonymous. The nomenclature of diseases:III. BMJ 1869;1(420) :55–57.

80. Philipson GH. On the Registration of Diseases. BMJ [Internet] 1869 Nov 6;2(462) :485–487. [doi: 10.1136/bmj.2.462.485]

81. Willshire WH. The Epidemic of Rubeola. Lancet [Internet] 1856 Jun;67(1710) :640. [doi: 10.1016/S0140-6736(02)55500-8]

82. Kesteven WB. The Epidemic of Rubeola. Lancet [Internet] 1856 Jun;67(1711) :671–672. [doi: 10.1016/S0140-6736(02)55536-7]

83. Tripe JW. The Rubeola Epidemic. Lancet [Internet] 1856 Jun;67(1713) :719–720. [doi: 10.1016/S0140-6736(02)59408-3]

84. Scattergood T. Morbilli and Rubeola. BMJ [Internet] 1870 Jan 29;1(474) :121–121. [doi: 10.1136/bmj.1.474.121]

85. Squire W. Remarks on Epidemic Roseola; Rosella, Rosalia, or Rubeola. BMJ [Internet] 1870 Jan 29;1(474) :99–100. [doi: 10.1136/bmj.1.474.99]

86. Forbes JA. Rubella: Historical Aspects. Arch Pediatr Adolesc Med [Internet] 1969 Jul 1;118(1) :5–11. [doi: 10.1001/archpedi.1969.02100040007002]

87. Ziring PR, Florman AL, Cooper LZ. The Diagnosis of Rubella. Pediatr Clin North Am [Internet] 1971 Feb;18(1) :87–97. [doi: 10.1016/S0031-3955(16)32524-X]

88. Banatvala JE, Brown D. Rubella. Lancet [Internet] 2004 Apr;363(9415) :1127–1137. [doi: 10.1016/S0140-6736(04)15897-2]

89. Murchison C. Clinical Lectures on Medicine. Lancet [Internet] 1870 Oct;96(2461) :595–598. [doi: 10.1016/S0140-6736(02)79936-4]

90. Lin Y, Michel J, Aiden EL, Orwant J, Brockman W, Petrov S. Syntactic Annotations for the Google Books Ngram Corpus. Proc ofthe 50th Annu Meet ofthe Assoc Comput Linguist Jeju, Republic of Korea: Association for Computational Linguistics,Stroudsburg, PA, USA; 2012. p. 169–174.

91. Putnam CP. Is German Measles (Rotheln or Rubella) an Independent Disease? Bost Med Surg J [Internet] 1893 Jul 13;129(2) :30–33. [doi: 10.1056/NEJM189307131290202]

92. Rota PA, Moss WJ, Takeda M, de Swart RL, Thompson KM, Goodson JL. Measles. Nat Rev Dis Prim [Internet] 2016 Dec 22;2(1) :16049. [doi: 10.1038/nrdp.2016.49]

93. Gastel B. Measles: A Potentially Finite History. J Hist Med Allied Sci [Internet] 1973;28(1) :34–44. [doi: 10.1093/jhmas/XXVIII.1.34]

94. Drutz JE. Measles: Its history and its eventual eradication. Semin Pediatr Infect Dis [Internet] 2001 Oct;12(4) :315–322. [doi: 10.1053/spid.2001.26640]

95. Rosen G. A history of public health. 1st ed. New York: M.D. Publications; 1958.

96. Panum PL. Observations Made During the Epidemic of Measles on the Faroe Islands in the Year 1846. Copenhagen: Bibliothek for Laeger; 1847.

97. Veale H. History of an Epidemic of Rötheln, with Observations on Its Pathology. Edinb Med J [Internet] 1866 Nov;12(5) :404–414. PMID:29646314

98. Tomkins H. The Diagnosis of Rötheln. BMJ [Internet] 1880 May 29;1(1013) :808–808. [doi: 10.1136/bmj.1.1013.808]

99. Treveris P. The Grete Herball: a tendril of ancient herbal traditions reaching into 16th century England. 1st ed. Treveris P, editor. London; 1526.

100. Encyclopaedia Britannica. 1st ed. Edinburgh, Scotland: A. Bell and C. Macfarquhar; 1771.

101. Paterson R. Observations on Corpora Lutea: Part I. Edinburgh Med Surg J [Internet] 1840 Jan 1;53(142) :49–67. PMID:30331001

102. Veale H. On the Etiology of Stricture of the Urethra. Edinb Med J [Internet] 1867 Jan;12(7) :602–606. PMID:29646509

103. Shulman ST. The History of Pediatric Infectious Diseases. Pediatr Res [Internet] 2004 Jan;55(1) :163–176. [doi: 10.1203/01.PDR.0000101756.93542.09]

104. Paterson R. An Account of the Rötheln of German Authors, Together with a Few Observations on the Disease as It Has Been Seen to Prevail in Leith and Its Neighbourhood. Edinburgh Med Surg J [Internet] 1840 Apr 1;53(143) :381–393. PMID:30330988

105. Maton WG. Some account of a rash liable to be mistaken for scarlatina. Med Trans Coll Physicians 1815;5:149–165.

106. Cooper LZ. The history and medical consequences of rubella. Rev Infect Dis 1985;7:2–10. PMID:3890105

107. Grimwood RM, Holmes EC, Geoghegan JL. A Novel Rubi-Like Virus in the Pacific Electric Ray (Tetronarce californica) Reveals the Complex Evolutionary History of the Matonaviridae. Viruses [Internet] 2021 Mar 31;13(4) :585. [doi: 10.3390/v13040585]

108. Ackerknecht EH. A short history of medicine. Baltimore, Maryland: Johns Hopkins University Press; 1982. ISBN:978-0-8018-2726-6

109. Neuburger M. Some relations between British and German medicine in the first half of the eighteenth century. Bull Hist Med 1945;17(3) :217–228.

110. Hess V, Mendelsohn JA. Sauvages’ Paperwork: How Disease Classification Arose from Scholarly Note-Taking. Early Sci Med [Internet] 2014 Nov 27;19(5) :471–503. [doi: 10.1163/15733823-00195p05]

111. International Congress of Medicine. Transactions of the International Congress of Medicine. Trans Int Congr Med London; 1881. p. 14–34.

112. International Medical Congress. Lancet [Internet] 1881 Mar;117(3001) :392–393. [doi: 10.1016/S0140-6736(02)33362-2]

113. Billings JS. International Medical Congress, London, 1881. Bost Med Surg J [Internet] 1881 Sep 8;105(10) :217–222. [doi: 10.1056/NEJM188109081051001]

114. International Medical Congress. Lancet [Internet] 1881 Mar;117(3003) :478. [doi: 10.1016/S0140-6736(02)13222-3]

115. The International Medical Congress. Nature [Internet] 1881 Aug;24(615) :338–339. [doi: 10.1038/024338a0]

116. The International Medical Congress. BMJ [Internet] 1881 Aug 13;2(1076) :300–305. [doi: 10.1136/bmj.2.1076.300]

117. The International Medical Congress. BMJ 1881;2(1082) :518–530. [doi: 10.1016/S0140-6736(01)93495-6]

118. International Medical Congress. BMJ [Internet] 1881 Oct 1;2(1083) :545–566. [doi: 10.1136/bmj.2.1083.545]

119. Barnett R. Rubella. Lancet [Internet] 2017 Apr;389(10077) :1386. [doi: 10.1016/S0140-6736(17)30890-5]

120. Warrack JS. The differential diagnosis of scarlet fever, measles, and rubella. BMJ [Internet] 1918 Nov 2;2(3018) :486–488. [doi: 10.1136/bmj.2.3018.486]

121. Editorial. Rubella in an Isolated Community. N Engl J Med [Internet] 1957 Feb 14;256(7) :323–324. [doi: 10.1056/NEJM195702142560712]

122. Gregg NM. Congenital Cataract Following German Measles in the Mother. Epidemiol Infect [Internet] 1991 Aug 15;107(1) :3–14. [doi: 10.1017/S0950268800048627]

123. Lambert N, Strebel P, Orenstein W, Icenogle J, Poland GA. Rubella. Lancet [Internet] 2015 Jun;385(9984) :2297–2307. [doi: 10.1016/S0140-6736(14)60539-0]

124. WHO. The Measles & Rubella Initiative supports implementation of the Measles & Rubella Strategic Framework 2021-2030 [Internet]. 2020. Available from: https://s3.amazonaws.com/wp-agility2/measles/wp-content/uploads/2020/11/measles_rubella_initiative_final_print.pdf

125. Zipf GK. Human Behavior and the Principle of Least Effort: An Introduction to Human Ecology. Cambridge, Massachusetts: Addison-Wesley Press; 1949. ISBN:4444006101174

126. Editorial. Reports of Meetings. N Engl J Med [Internet] 1941 Jan 30;224(5) :216–217. [doi: 10.1056/NEJM194101302240517]

127. Editorial. Miscellany. N Engl J Med [Internet] 1931 Apr 30;204(18):913–947. [doi: 10.1056/NEJM193104302041806]

## References

Jobin, A., M. Ienca, and E. Vayena. 2019. The global landscape of AI ethics guidelines. Nature Machine Intelligence 1(9) :389–399. doi: 10.1038/s42256-019-0088-2.

Page, M. J., D. Moher, P. M. Bossuyt, I. Boutron, T. C. Hoffmann, C. D. Mulrow, L. Shamseer, J. M. Tetzlaff, E. A. Akl, S. E. Brennan, R. Chou, J. Glanville, J. M. Grimshaw, A. Hróbjartsson, M. M. Lalu, T. Li, E. W. Loder, E. Mayo-Wilson, S. McDonald, L. A. McGuinness, L. A. Stewart, J. Thomas, A. C. Tricco, V. A. Welch, P. Whiting, and J. E. McKenzie. 2021. PRISMA 2020 explanation and elaboration: updated guidance and exemplars for reporting systematic reviews. BMJ: n160. doi: 10.1136/bmj.n160.

PRISMA. 2015. Transparent reporting of systematic reviews and meta-analyses: PRISMA. http://www.prisma-statement.org/

Zeraatkar, K., and M. Ahmadi. 2018. Trends of infodemiology studies: a scoping review. Health Information & Libraries Journal 35(2) :91–120. doi: 10.1111/hir.12216.

## References

1. Enserink M. Amid Heightened Concerns, New Name for Novel Coronavirus Emerges. Science [Internet] 2013 May 10;340(6133) :673–673. PMID:23661733

2. Dowell SF, Bresee JS. Pandemic lessons from Iceland. Proc Natl Acad Sci [Internet] 2008 Jan 29;105(4) :1109–1110. [doi: 10.1073/pnas.0711535105]

3. Hoppe T. “Spanish Flu”: When Infectious Disease Names Blur Origins and Stigmatize Those Infected. Am J Public Health [Internet] 2018 Nov;108(11) :1462–1464. [doi: 10.2105/AJPH.2018.304645]

4. Have we had Hong Kong flu before? Lancet [Internet] 1969 May;293(7601) :929–930. [doi: 10.1016/S0140-6736(69)92556-2]

5. Peckham R. Viral surveillance and the 1968 Hong Kong flu pandemic. J Glob Hist [Internet] 2020 Nov 6;15(3) :444–458. [doi: 10.1017/S1740022820000224]

6. Chang, BS C, Salerno, MSc A, Hsu, MD, mph EB. Perspectives on xenophobia during epidemics and implications for emergency management. J Emerg Manag [Internet] 2020 Dec 10;18(7) :23–29. [doi: 10.5055/jem.2020.0521]

7. Wexler A. Stigma, history, and Huntington’s disease. Lancet [Internet] 2010 Jul;376(9734) :18–19. [doi: 10.1016/S0140-6736(10)60957-9]

8. Rawlins M. Huntington’s disease out of the closet? Lancet [Internet] 2010 Oct;376(9750) :1372–1373. [doi: 10.1016/S0140-6736(10)60974-9]

9. Spinney L. Uncovering the true prevalence of Huntington’s disease. Lancet Neurol [Internet] 2010 Aug;9(8) :760–761. [doi: 10.1016/S1474-4422(10)70160-5]

10. Editorial. Dispelling the stigma of Huntington’s disease. Lancet Neurol [Internet] 2010 Aug;9(8) :751. [doi: 10.1016/S1474-4422(10)70170-8]

11. Hu Z, Yang Z, Li Q, Zhang A. The COVID-19 Infodemic: Infodemiology Study Analyzing Stigmatizing Search Terms. J Med Internet Res [Internet] 2020 Nov 16;22(11) :e22639. PMID:33156807

